## Supplementary material for "The LINC complex transmits integrin-dependent tension to the nuclear lamina and represses epidermal differentiation": All Figure Supplements

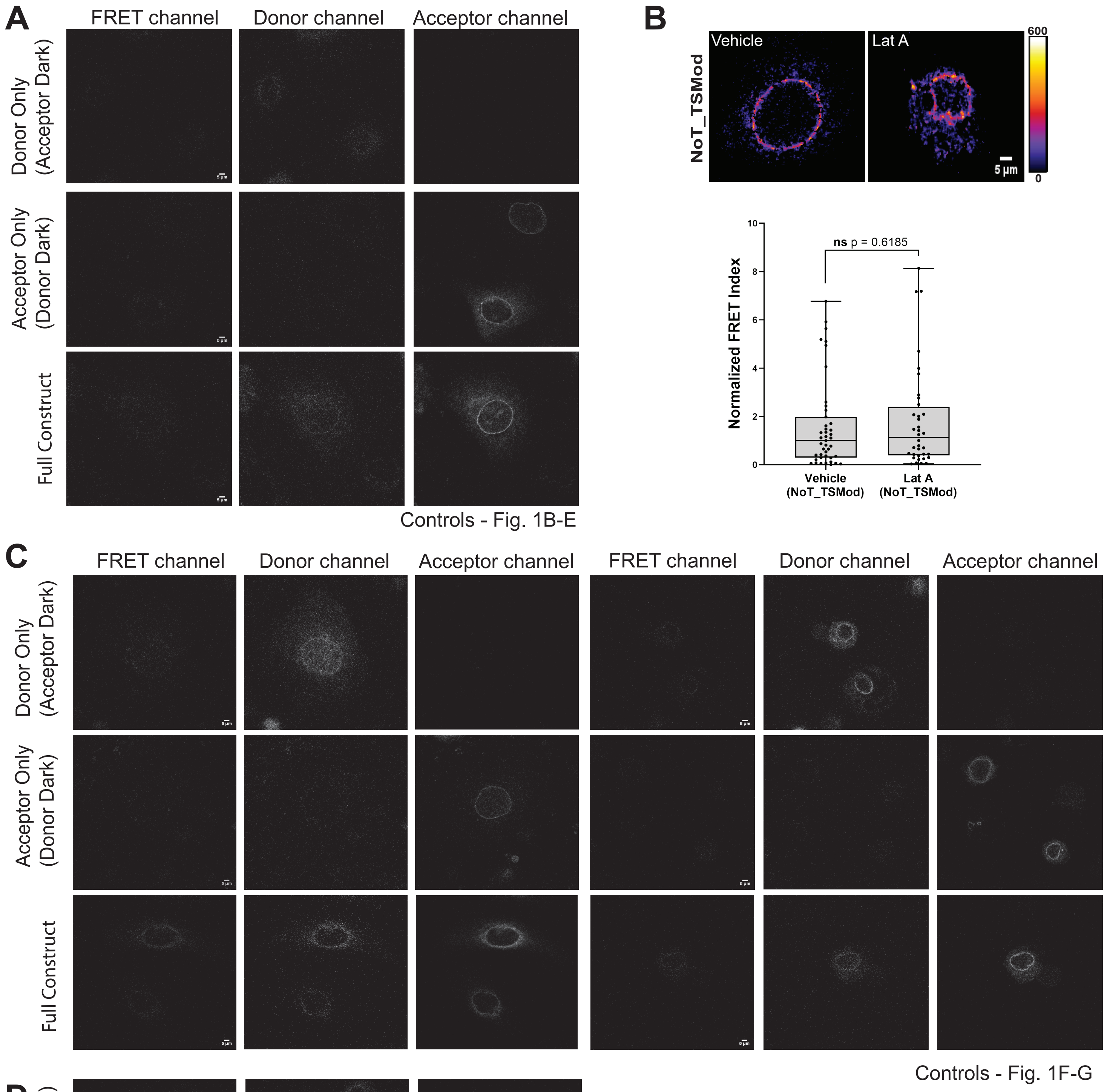


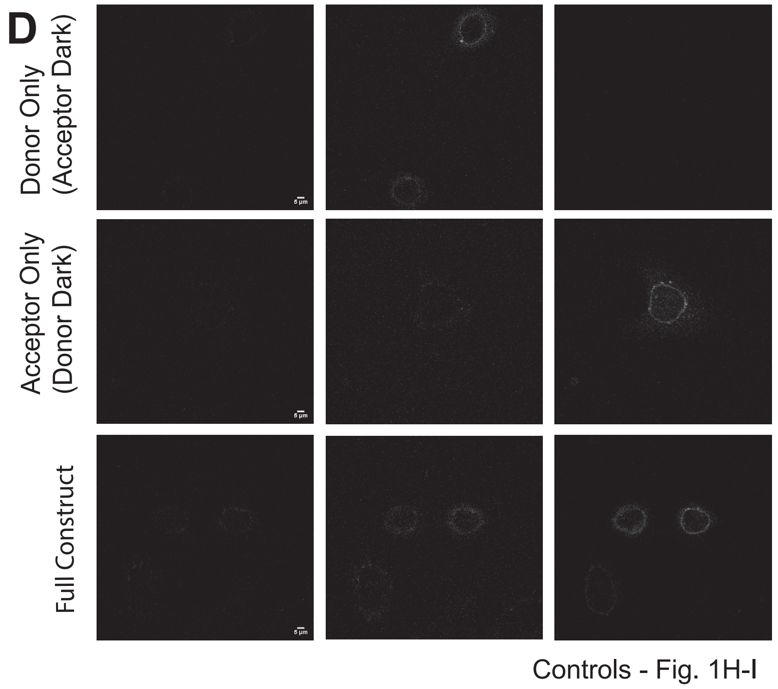


### Figure 1- figure supplement 1

**Validation controls for the N2G-JM-TSMod**

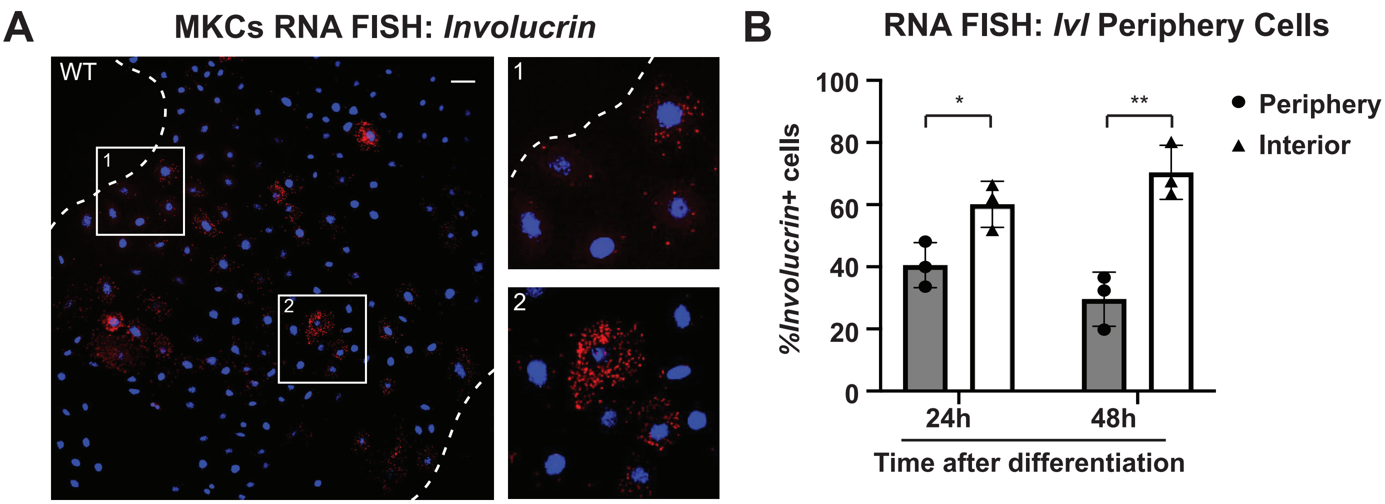


### Figure 2-figure supplement 1

**The differentiation marker involucrin (*Ivl*) is expressed at higher levels in the colony interior than at the colony periphery.**

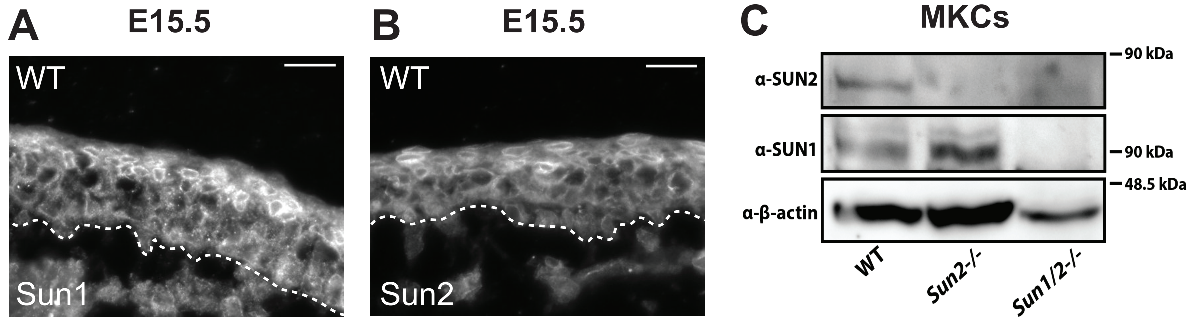


### Figure 3-figure supplement 1

**SUN1 and SUN2 are expressed throughout the epidermis.**

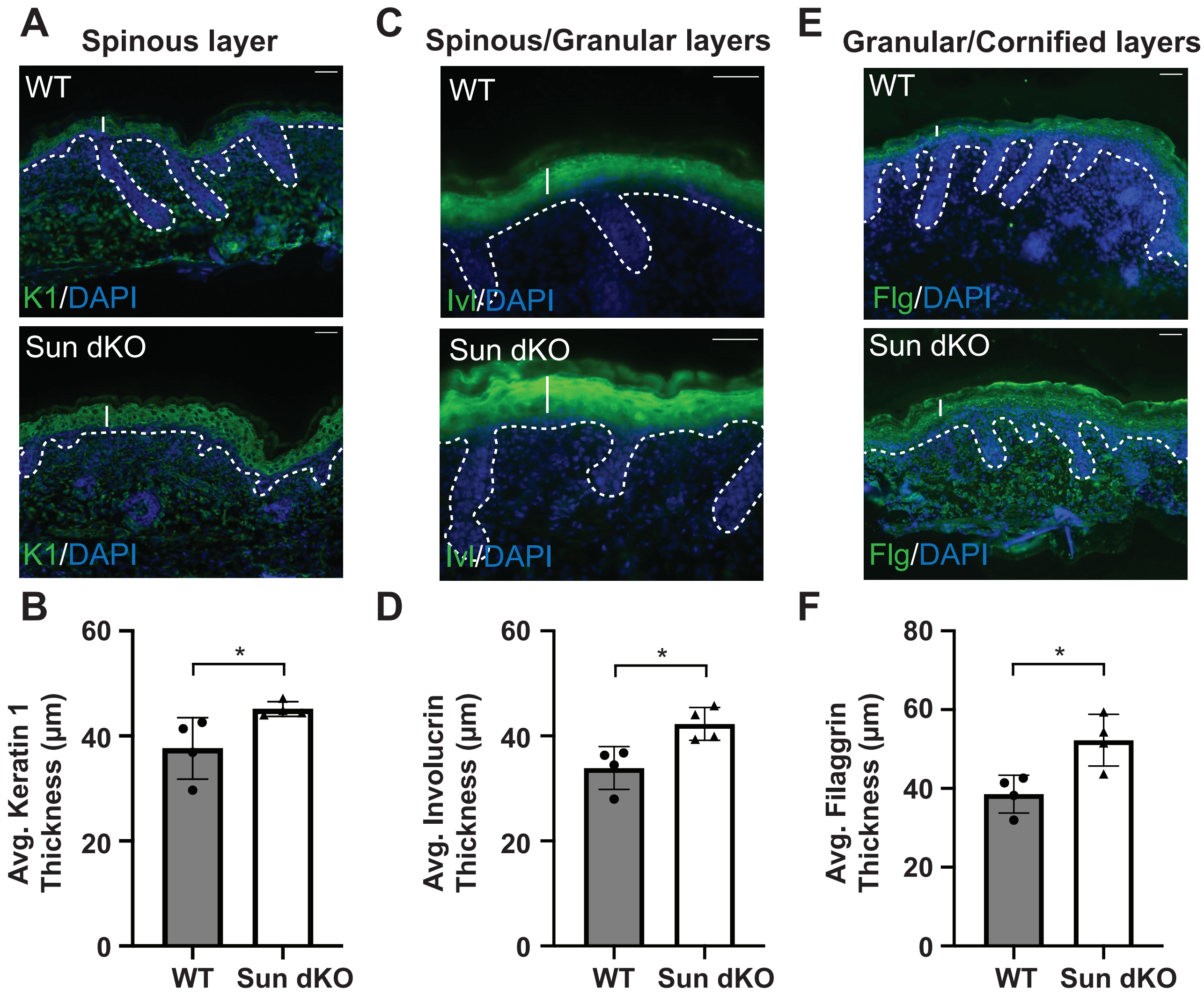


#### Figure 3-figure supplement 2

**Thickening of the differentiated layers in the Sun dKO epidermis.**

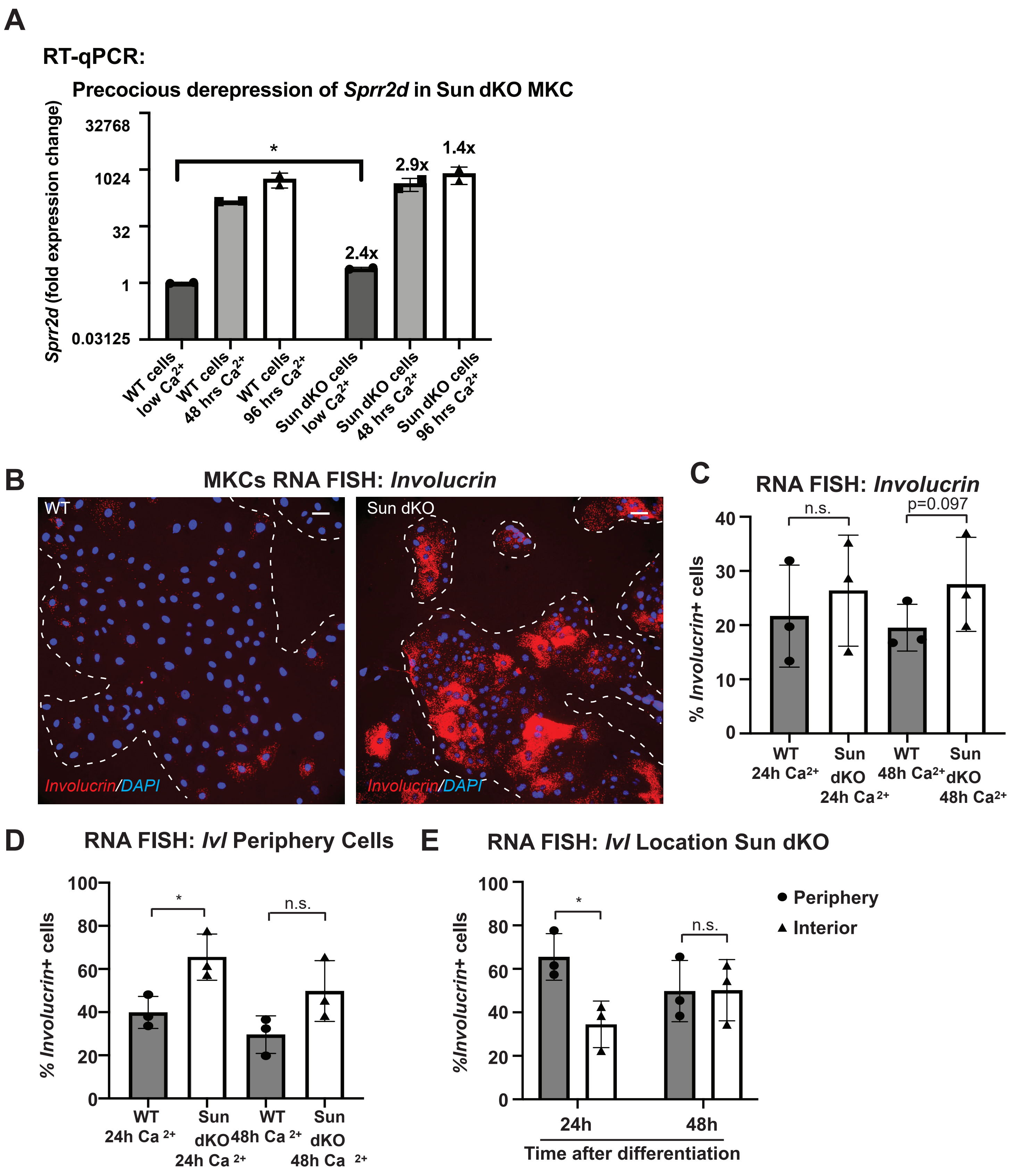


#### Figure 4-figure supplement 1

**Additional evidence for precocious differentiation in Sun dKO MKCs.**

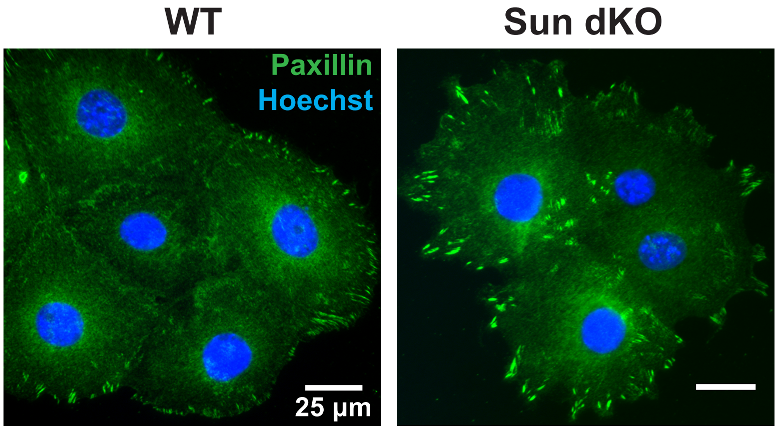


#### Figure 4-figure supplement 2

**Cohesive Sun dKO MKC colonies have focal adhesions at the colony periphery.**

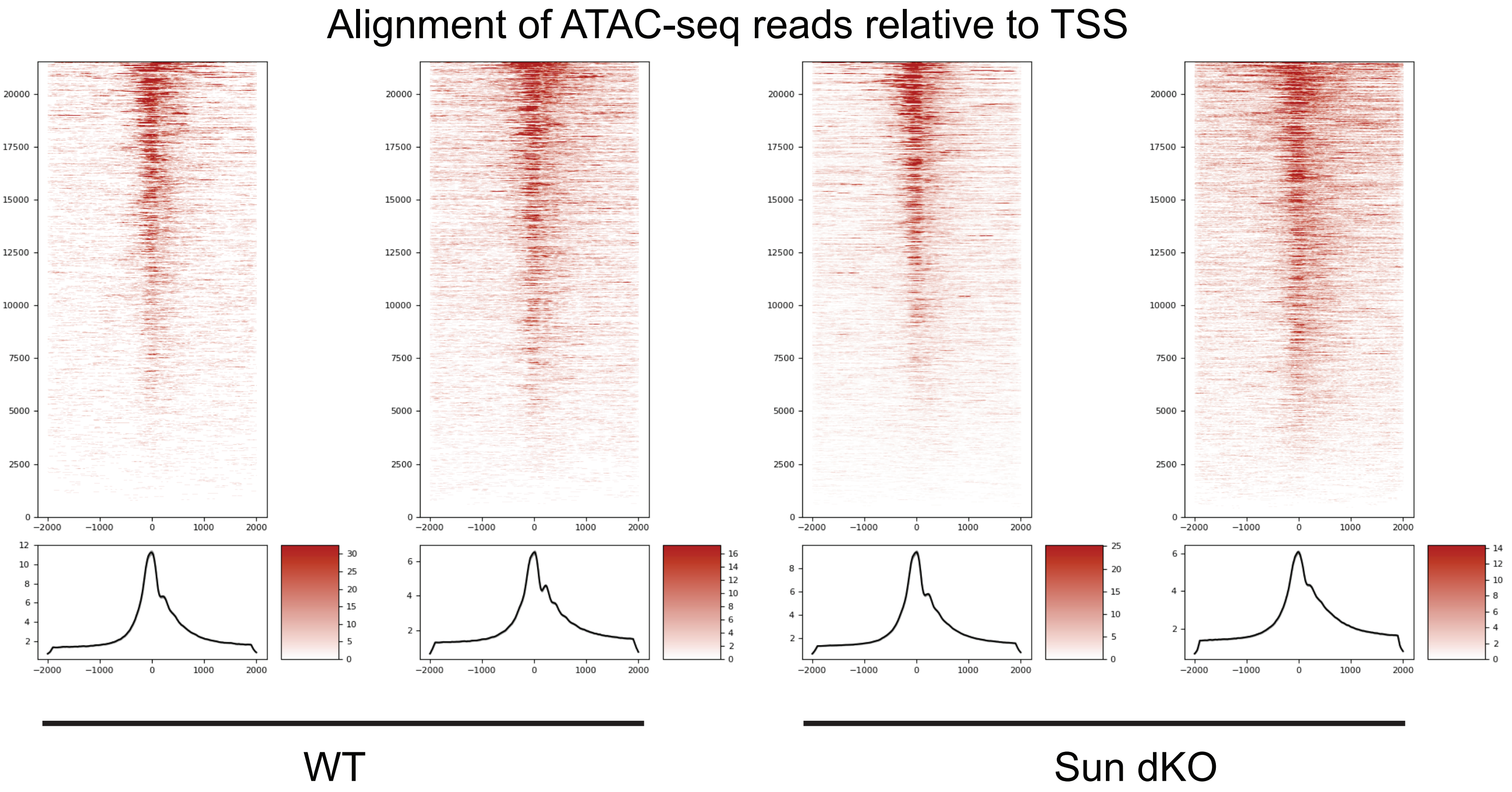


**Figure 5 Supplement 1**

**Quality assessment for each replicate of ATAC-seq data.**

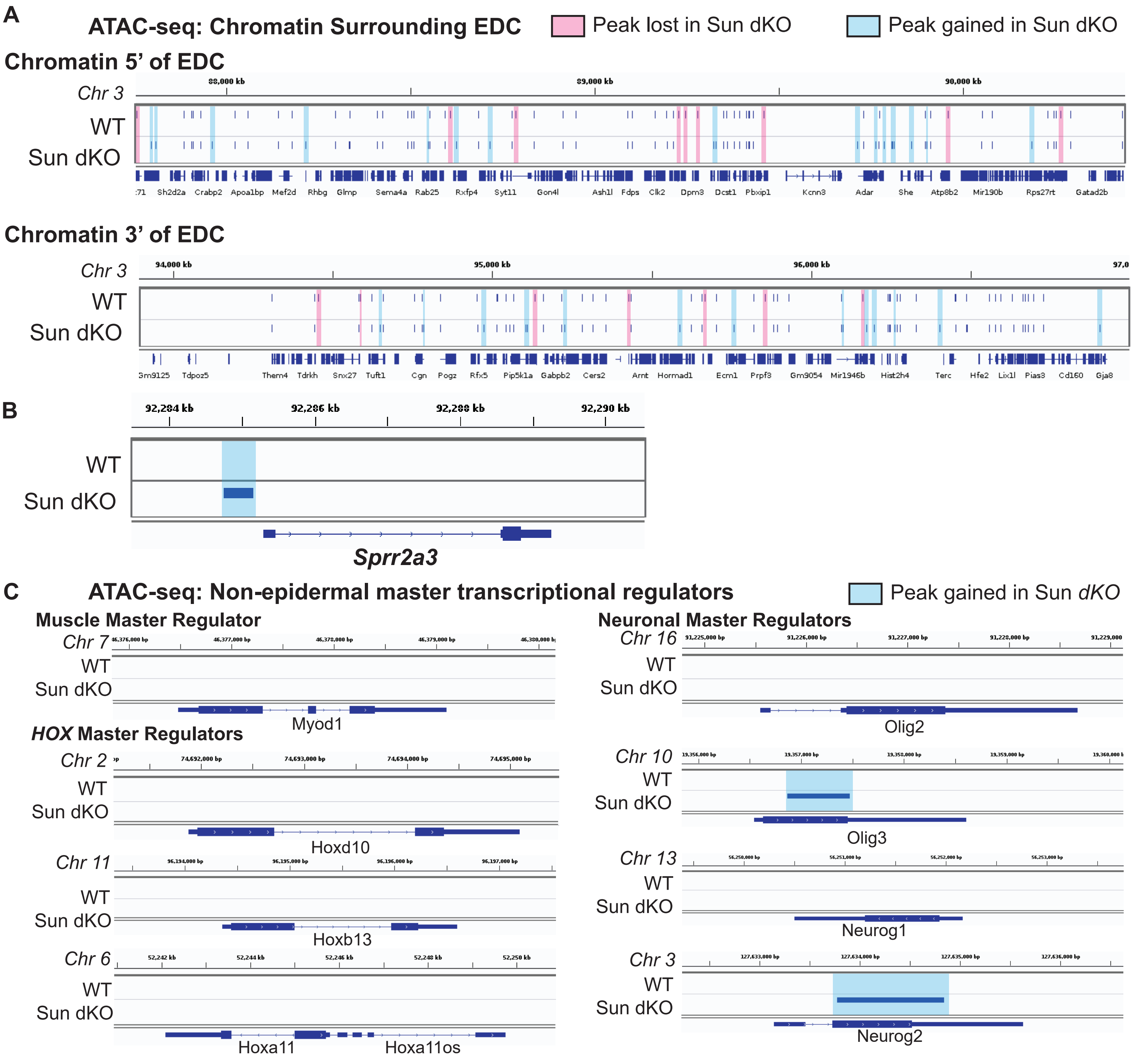


### Figure 5-figure supplement 2

**Increased accessibility within the EDC is specific as assessed by ATAC-seq in Sun dKO MKCs cultured in low calcium media.**
